# Early adjunct anti-PD-L1 immunotherapy improves outcomes and restores infection-induced immune paralysis in mice with invasive pulmonary mucormycosis

**DOI:** 10.1101/2025.05.06.652431

**Authors:** Sebastian Wurster, Jezreel Pantaleon Garcia, Yongxing Wang, Nathaniel D. Albert, Uddalak Bharadwaj, Lauren Matula, Russell E. Lewis, Scott E. Evans, Dimitrios P. Kontoyiannis

## Abstract

Invasive pulmonary mucormycosis (IPM) is a severe opportunistic mold infection whose outcome is predominantly host driven. Preclinical proof-of-concept studies and clinical case reports in salvage therapy settings suggested a benefit of immune checkpoint inhibitors (ICIs) in IPM management. However, the kinetics of infection-induced immune paralysis and optimal timing of ICI therapy remain poorly understood. Here, we performed sequential nCounter-based transcriptomics on lung tissue of cyclophosphamide-immunosuppressed mice with IPM (*Rhizopus arrhizus* infection) to dynamically study the pulmonary immune environment. Within 7 days after infection, lungs of mice with IPM showed reversal of early proinflammatory signaling, impaired T-cell signaling, and upregulation of exhaustion markers. Similar immune paralysis signatures were seen in mice with invasive pulmonary aspergillosis and fusariosis. For therapeutic studies, Mucorales-active antifungal therapy with isavuconazonium sulfate (ISAV) was initiated on day 3 after IPM infection, along with anti-PD-L1 or a non-targeting isotype antibody (control) given either on days 3+5 (early) or days 6+8 (late). Both early and late adjunct anti-PD-L1 therapy were well-tolerated and significantly improved morbidity/mortality outcomes compared to ISAV + isotype. Notably, early adjunct anti-PD-L1 therapy promoted significantly stronger innate immune cell activation, upregulation of key cytokine pathways, reinvigoration of T-helper-cell signaling, and reversal of IPM-induced exhaustion signals than both late adjunct anti-PD-L1 and isotype control. These findings indicate that combined antifungal and early immunomodulatory therapy may be an important strategy to intercept immune paralysis and improve outcomes in immunocompromised hosts with IPM, inviting further preclinical and clinical exploration of early host-directed interventions to treat deadly mold pneumonias.

## Introduction

Mucormycosis is an aggressive opportunistic mold infection affecting a broad spectrum of susceptible hosts including patients with hematologic malignancies, high-dose glucocorticosteroid use, uncontrolled diabetes mellitus, COVID-19, or severe trauma^1,2^. Its most common manifestation in cancer patients receiving cytotoxic chemotherapy is invasive pulmonary mucormycosis (IPM)^1,2^. Even with adequate antifungal therapy, IPM has a poor prognosis, with an overall mortality of 50% that approaches 100% in patients with disseminated disease or persistent neutropenia^3,4^.

Because the efficacy of conventional antifungal therapy is limited and mitigation of the underlying breaches to antifungal host defense is the preeminent prognostic determinant in patients with IPM^4^, adjunct immune enhancement strategies hold significant promise to improve outcomes. Specifically, there is growing interest in the use of readily available and rapidly deployable non-cellular immune enhancers to bolster host defense against pathogenic molds^5^, including immune checkpoint inhibitors (ICIs)^6,7^.

Checkpoint pathways, such as the Programmed Cell Death Protein 1 (PD-1) pathway, form part of a tightly controlled network of stimulatory and regulatory signals governing human immune homeostasis^8^. While critical to mediate immunologic self-tolerance and prevent overzealous inflammatory responses, checkpoint pathways can also drive immune exhaustion in settings of persistent inflammation and antigen exposure due to malignancies or infections^7,8^. Thus, ICI antibodies targeting these pathways have revolutionized modern cancer medicine^9^ and are increasingly studied as an adjunct immunotherapy against various infections diseases^10^, including opportunistic mold infections^7^.

Corroborating anecdotal clinical reports of ICIs used as salvage therapy for otherwise intractable mold infections^11–14^, we and others have previously shown significant benefits of ICI therapy in mouse models of various invasive fungal diseases (IFDs)^7,15–19^. Specifically, we have demonstrated significantly improved morbidity, mortality, and fungal clearance after low-dose (mono-)therapy with antibodies blocking PD-1 or its main ligand Programmed Death Ligand 1 (PD-L1) in neutropenic mice with IPM^18^. However, optimal deployment and timing of ICIs for multimodal therapy of IPM remains elusive^7,18^.

We hypothesized that pulmonary mold infections are associated with a rapid shift of host immunity from initial infection-induced inflammation to paralyzed host defense that might be a key “doorway” to fungal dissemination. Therefore, we herein dynamically study the host response to IPM and other mold pneumonias in mice with chemotherapy-induced neutropenia. Furthermore, we compare the benefit of early versus late adjunct ICI therapy combined with first-line antifungal therapy in mice with IPM. We show that IPM and other deadly mold infections trigger rapidly emerging immune paralysis and induction of checkpoint pathways. We further demonstrate that IPM-induced immunopathology is strongly attenuated upon multimodal therapy with antifungals and early adjunct anti-PD-L1 (α-PD-L1), resulting in significantly improved morbidity/mortality outcomes and enhanced fungal clearance.

## Results

### IPM infection induces severe and rapidly emerging immune paralysis

To study pulmonary immune dynamics, we used a murine IPM infection model with cyclophosphamide-induced neutropenia and intranasal inoculation with *Rhizopus arrhizus*, the most common cause of IPM^20^ (**Fig. 1A**). Untreated mice with IPM developed severe and rapidly evolving distress, with median morbidity/mortality scores of 1.5 on day 4, 1.8 on day 7, and 2.9 on day 10 (modified murine sepsis score [MSS]; 0 = healthy – 3 = moribund; 4 = death before the time of assessment)^18,21^.

**Figure 1.**
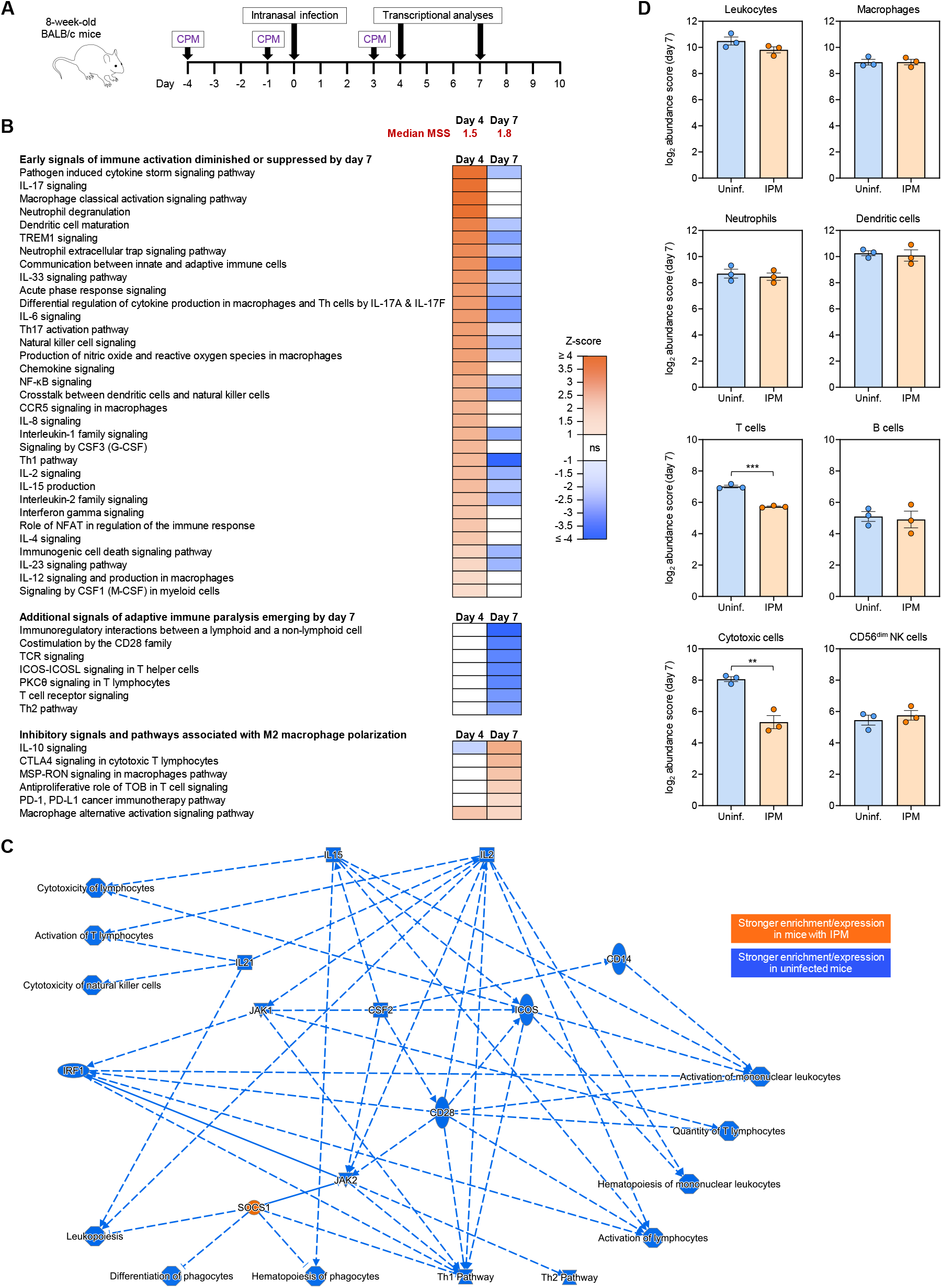
The pulmonary immune environment shifts from immune activation to a state of immune paralysis within one week of IPM infection. (**A**) Timeline of experimental procedures. (**B**) nCounter-based canonical pathway enrichment analyses performed on lung tissue of CPM-immunosuppressed mice with IPM on days 4 and 7 after infection. Data is compared to uninfected CPM-immunosuppressed mice. Selected immune-related pathways with an absolute z-score ≥1 and Benjamini Hochberg-adjusted p-value <0.05 for at least one of the two timepoints are shown. Orange and blue color indicate significantly greater enrichment in mice with IPM and uninfected mice, respectively. (**C**) Network of changes to the pulmonary immune transcriptome by day 7 after IPM infection compared to uninfected CPM-immunosuppressed mice, as predicted by Ingenuity Pathway Analysis. (**D**) Log_2_-transformed abundance scores of immune cell subsets determined by nCounter-based cell type profiling. Individual datapoints, means, and standard errors of the mean are shown. Unpaired two-sided t-test. ** p<0.01, *** p<0.001. (**B-D**) All analyses are based on n = 3 mice per group and timepoint. Abbreviations (not including gene symbols in panel **D**): CCR = C-C-type complement receptor, CD = cluster of differentiation, CPM = cyclophosphamide, CTLA-4 = cytotoxic T lymphocyte-associated protein 4, (G/M)-CSF = (granulocyte/macrophage) colony stimulating factor, ICOS(L) = inducible T-cell co-stimulator (ligand), IL = interleukin, IPM = invasive pulmonary mucormycosis, MSP-RON = macrophage stimulating protein & *recepteur d’origine nantais*, MSS = (modified) murine sepsis score, NFAT = nuclear factor of activated T cells, NF-κB = nuclear factor kappa B, ns = not significant, PD-1 = programmed cell death protein 1, PD-L1 = programmed death ligand 1, PKC = protein kinase C, TCR = T-cell receptor, Th = T-helper (cell), TOB = Transducer of ERBB2, TREM = triggering receptors expressed on myeloid cells.

We then performed nCounter-based transcriptomics and pathway enrichment analysis on lung tissue obtained on days 4 and 7 post-infection. Expectedly, mice with IPM displayed strong early innate immune activation compared to uninfected immunosuppressed mice, including upregulation of several inflammatory cytokine pathways (e.g., IL-6 and IL-8 signaling), activation of neutrophil effector pathways, classical (M1) macrophage and natural killer (NK)-cell signaling, and enhanced intercellular crosstalk (**Fig. 1B**). Likewise, several pathways associated with induction of type-1 (Th1) and type-17 (Th17) T-helper-cell responses were upregulated in mice with IPM (**Fig. 1B**). However, already by day 4 post-infection, several genes associated with T-cell activation (e.g., *ICOS*), NK-cell activation (e.g., *CD160*), and cytotoxicity (e.g., *GZMA, GZMB*) were significantly suppressed in mice with IPM compared to uninfected mice (**Fig. S1**).

By day 7 post-infection, proinflammatory responses and innate immune activation had markedly decreased, despite increasing IPM severity. Residual inflammatory signals were counter-balanced by signs of impaired intercellular crosstalk and immune paralysis (**Fig. 1B**). Specifically, mice with IPM displayed strong suppression of T-cell signaling, attenuation of several critical cytokine pathways (e.g., IL-2 and IL-6), and impaired innate immune effector responses (e.g., suppressed NK-cell and neutrophil signaling) compared to uninfected immunosuppressed controls. Moreover, mice with IPM showed upregulation of immunoregulatory cytokines (IL-10), unfavorable polarization of macrophages toward an alternative (M2) phenotype (e.g., MSP-RON signaling^22^), and induction of immune checkpoint pathways, including the PD-1/PD-L1 and CTLA4 pathways (**Fig. 1B-C**).

Paralysis of antifungal immunity by day 7 post-infection was further corroborated by transcriptional cell abundance scoring, indicating significantly reduced enrichment of transcripts attributed to T cells (log_2_ abundance score 5.7 vs. 7.0, p<0.001) and “cytotoxic cells” (log_2_ abundance score 5.3 vs. 8.1, p=0.004) in mice with IPM compared to uninfected controls (**Fig. 1D**). Collectively, these transcriptional signals indicate a rapid shift from immune activation toward severe immune paralysis within one week of IPM infection.

### Rapidly emerging immune paralysis is a common hallmark of IPM and other mold pneumonias

Next, we sought to test whether these immune paralysis signals are unique to IPM or also seen in other common opportunistic mold pneumonias. Therefore, we performed nCounter-based transcriptomics on lung tissue from mice with invasive pulmonary aspergillosis (IPA; *Aspergillus fumigatus*) and fusariosis (IF; *Fusarium solani*) using pre-optimized inoculums that yielded morbidity (MSS) scores comparable to those in mice with IPM (**Fig. S2A**). While only few upregulated transcripts were found on day 7 post-infection, especially in mice with IPM and IPA, all three mold pneumonias led to significant downregulation of immune-related transcripts, suggesting a predominantly immunosuppressive environment (**Fig. 2A**). Notably, a significant proportion of downregulated transcripts (n = 25) were shared among all three mold pneumonias (**Fig 2B**). These transcripts were mostly related to chemokine signaling, T- and NK-cell activation, and cytotoxic effector responses (box in **Fig 2B**).

**Figure 2.**
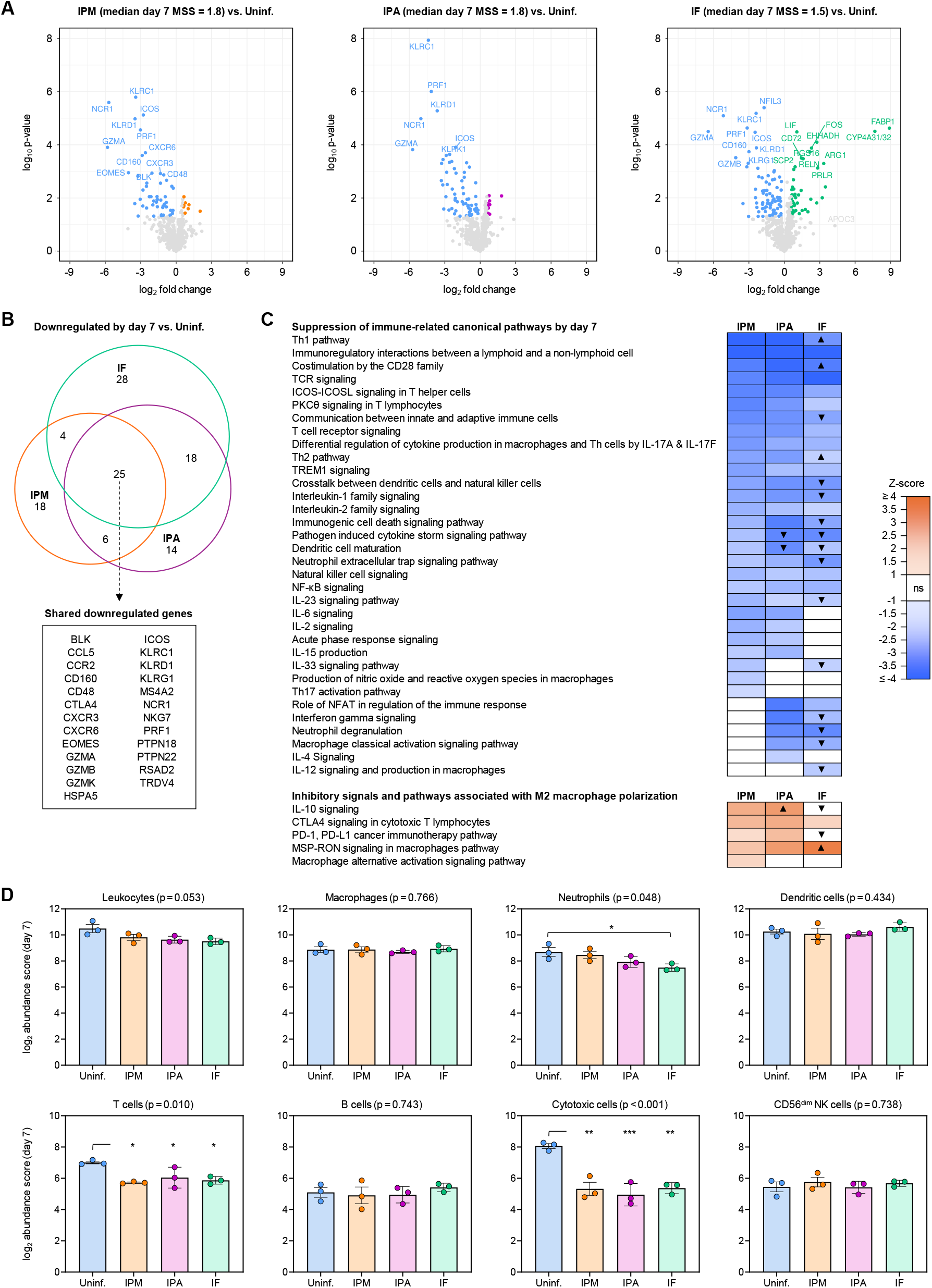
Key features of immune paralysis are shared among common mold pneumonias affecting immunocompromised hosts. (**A**) Volcano plots comparing gene expression levels in lung tissue of mice with IPM, IPA, and IF versus uninfected mice. Analyses are based on the 785-gene nCounter Immune Exhaustion panel performed on day 7 post-infection. Significantly differentially expressed genes (absolute fold change >1.5, p<0.05) are highlighted. (**B**) Euler diagram summarizing numbers of unique and shared downregulated genes in infected mice according to the type of mold pneumonia. (**C**) Canonical pathway enrichment in mice with IPM, IPA, and IF compared to uninfected mice. Selected immune-related pathways with an absolute z-score ≥1 and Benjamini Hochberg-adjusted p-value <0.05 for at least one of the mold pneumonias are shown. Orange and blue color indicate significantly greater enrichment in infected and uninfected mice, respectively. Arrows in the IPA and IF columns indicate significantly increased (▴) or decreased (▾) enrichment compared to mice with IPM. See **Suppl. Datasheet** for complete data. (**D**) Log_2_-transformed abundance scores of immune cell subsets determined by nCounter-based cell type profiling. Individual datapoints, means, and standard errors of the mean are shown. One-way analysis of variance (p-values above panels) with Tukey’s post-test for pairwise comparisons (asterisks). * p<0.05, ** p<0.01, *** p<0.001. (**A-D**) All analyses are based on n = 3 mice per group and timepoint. Abbreviations (not including gene symbols in panels **A** and **B**): CD = cluster of differentiation, CTLA-4 = cytotoxic T lymphocyte-associated protein 4, ICOS(L) = inducible T-cell co-stimulator (ligand), IL = interleukin, IF = invasive fusariosis, IPA = invasive pulmonary aspergillosis, IPM = invasive pulmonary mucormycosis, MSS = (modified) murine sepsis score, NFAT = nuclear factor of activated T cells, NF-κB = nuclear factor kappa B, ns = not significant, PD-1 = programmed cell death protein 1, PD-L1 = programmed death ligand 1, PKC = protein kinase C, TCR = T-cell receptor, Th = T-helper (cell), TREM = triggering receptors expressed on myeloid cells.

We then compared differential pathway enrichment according to the type of mold pneumonia (**Fig. 2C**). Differences in pathway enrichment were minimal between mice with IPA and IPM, especially with regards to impaired T-cell signaling and upregulation of exhaustion markers. Modest differences were noted between IPM and IF, with the latter being associated with stronger impairment of innate immune cell and cytokine signaling but less pronounced T-cell paralysis than IPM (arrowheads in **Fig. 2C**).

Despite these granular differences, 37 out of 46 immune-related pathways (80%) that were significantly suppressed on day 7 after IPM infection were shared with at least one of the other mold pneumonias; 28 (61%) were shared among all three infections (**Suppl. Datasheet**). Specifically, pathways related to T-cell activation and Th1 signaling were among the top-ranked downregulated pathways in all three mold pneumonias when compared to uninfected immunosuppressed animals (**Fig. 2C, Fig. S2B**). These findings indicate that rapidly emerging infection-induced immune paralysis is a common immunopathogenic hallmark of several clinically important mold pneumonias.

This conclusion was further supported by transcriptional cell abundance scoring revealing a consistently reduced abundance of transcripts related to T cells (p=0.010) and “cytotoxic cells” (p<0.001) in all three mold pneumonias, whereas only IF led to a reduced abundance of neutrophil-associated transcripts (**Fig. 2D**).

### Early adjunct immunotherapy with α-PD-L1 is well tolerated and impactfully improves outcomes in mice with IPM

Focusing on *Rhizopus* infection, our primary pathogen of interest, we then compared therapeutic outcomes in mice receiving Mucorales-active antifungal therapy with isavuconazonium sulfate (ISAV; 64 mg/kg twice daily by oral gavage) in combination with a previously established sub-oncologic intraperitoneal dose of 0.25 mg/kg α-PD-L1 or the corresponding non-targeting isotype antibody as control (**Fig. 3A**)^17,18^. Adjunct immunotherapy was given either on days 3+5 (early) or days 6+8 (late). This setup allowed us to compare the benefits of intercepting immune exhaustion versus mitigating immune paralysis signals after they had manifested (**Fig. 1B**). Additionally, our timelines ensured that therapeutic interventions were only initiated once the mice showed robust signs of infection-induced distress (MSS >1.0).

**Figure 3.**
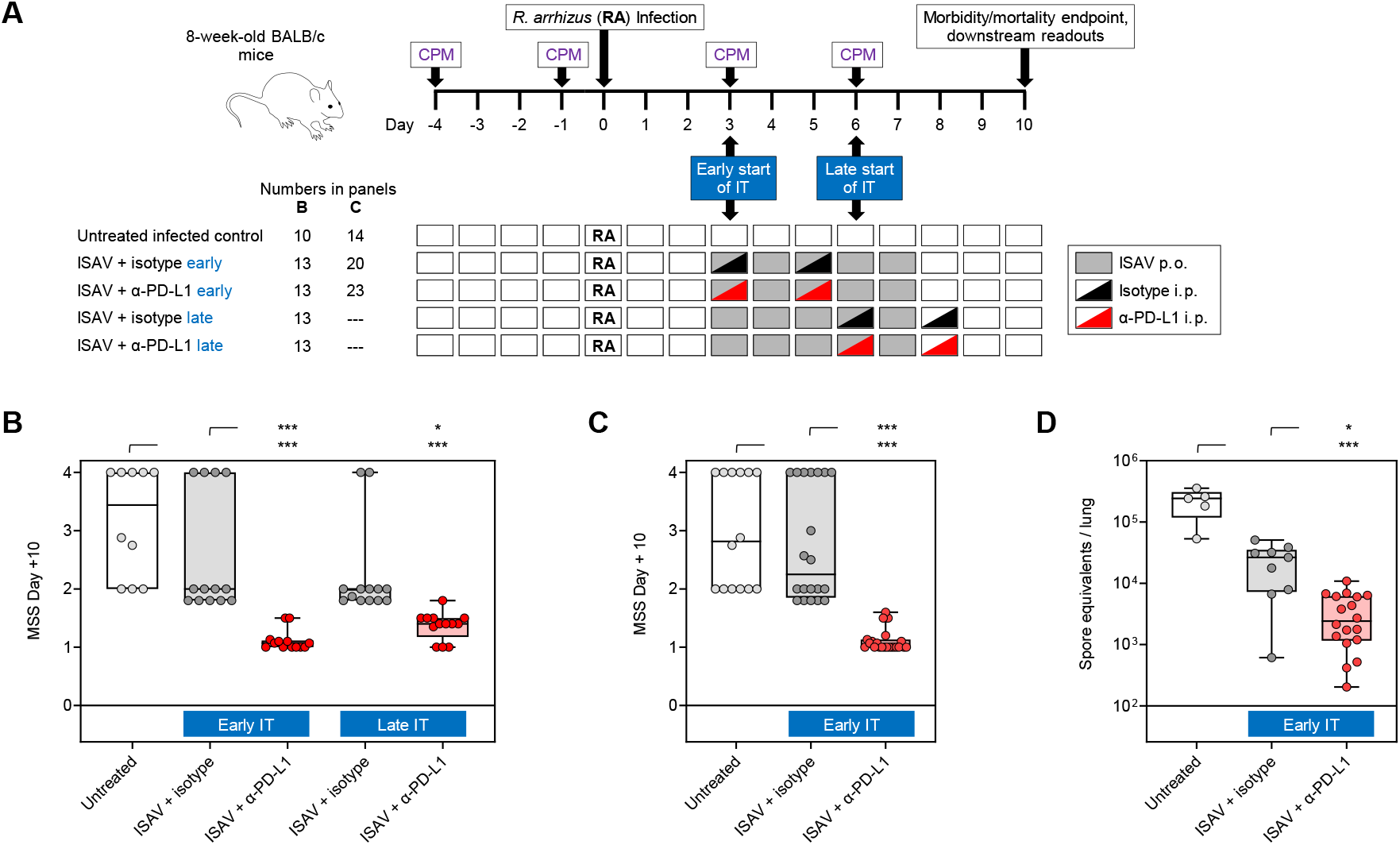
Combination therapy with ISAV and early adjunct α-PD-L1 significantly improves morbidity, mortality, and fungal clearance in mice with IPM. (**A**) Timeline of experimental procedures. (**B-C**) MSS scores on day 10 post-infection according to the treatment arm considering (**B**) only experiments that included all five groups (N = 10-13 mice per group, 2 independent experiments) or (**C**) all available datapoints for early adjunct immunotherapy (N = 14-23 mice per group, 4 independent experiments). (**D**) Fungal burden in lung tissue homogenates of mice that survived until day 10 in non-moribund condition. N = 5-18 mice per group, 3 independent experiments. (**B-D**) Individual data points, quartiles (boxes), and ranges (whiskers) are shown. Kruskal-Wallis test with Dunn’s post-test. * p<0.05, *** p<0.001. Abbreviations: α = anti, CPM = cyclophosphamide, IPM = invasive pulmonary mucormycosis, ISAV = isavuconazonium sulfate, IT = immunotherapy, MSS = (modified) murine sepsis score, PD-L1 = programmed death ligand 1, RA = *Rhizopus arrhizus*.

Untreated infected mice developed severe IPM, with 50% 10-day mortality and a median day-10 MSS of 3.4. Antifungal therapy with ISAV plus mock immunotherapy with the isotype antibody reduced 10-day mortality compared to untreated infected mice (15-31%). However, considerable morbidity persisted even with ISAV therapy (plus isotype), with a median day-10 MSS of 2.0 for both early and late isotype control groups. In contrast, early adjunct α-PD-L1 therapy initiated concomitantly with ISAV led to universal survival and strongly improved day-10 MSS (median, 1.1; p<0.001 versus ISAV + isotype). Although statistically significant, the adjunct therapeutic benefit of α-PD-L1 was less pronounced (median day-10 MSS, 1.4) when initiated late (days 6+8) (**Fig. 3B**). The strong benefit of early adjunct α-PD-L1 therapy was further corroborated when including morbidity/mortality data from additional replicate experiments performed for biospecimen sampling to facilitate downstream readouts (**Fig. 3C**). Moreover, sequential MSS scoring in a representative experiment confirmed rapid attenuation of morbidity in mice with IPM receiving ISAV plus early adjunct α-PD-L1 therapy (**Fig. S3**).

As a secondary efficacy endpoint, we assessed fungal burden in mice surviving until day 10 post-infection in non-moribund condition. Untreated mice with IPM showed substantial fungal burden, with a median of 2.4×10^5^ spore equivalents per lung (**Fig. 3D**). ISAV therapy plus isotype led to a 1-log_10_ reduction in median fungal burden (2.6×10^4^ spore equivalents per lung) and a further 1-log_10_ reduction when adding early adjunct α-PD-L1 (p=0.021), with a median of 2.4×10^3^ spore equivalents per lung (p<0.001 compared to untreated infected mice; **Fig. 3D**).

Immunotoxicity has been a common concern about the use of potent immunomodulators, such as ICIs, for infection management^7,10^. In our study, adjunct α-PD-L1 at sub-oncologic dosing was well-tolerated, with no adverse reactions or evident immunotoxicity after the injections. To further preclude systemic immunotoxicity, we performed a multiplexed cytokine/chemokine assay on serum from mice with IPM that received ISAV plus either early adjunct α-PD-L1 or isotype. Serum levels of all tested cytokines and chemokines were very low in either group on both days 7 and 10 post-infection, with IL-4 being the only cytokine with median levels >10 pg/mL (**Table S1**). No significant elevations in serum cytokine/chemokine levels were detected after adjunct α-PD-L1 therapy (**Table S1**). Likewise, no signs of pneumonitis, were seen on lung tissue sections (**Fig. S4**).

### Early adjunct α-PD-L1 therapy restores IPM-induced immune paralysis and enhances innate and adaptive antifungal immunity

On lung tissue transcriptomics, antifungal therapy with ISAV (plus isotype) delayed the tipping point from immune activation to full-blown immune paralysis. However, the mice displayed a “tolerogenic” immune state, with signs of beginning impairment of classical (M1) macrophage and NK-cell signaling (**Fig. S5**). In contrast, combination of ISAV with early adjunct α-PD-L1 significantly attenuated signals of adaptive immune exhaustion and restored most innate effector pathways compared to both ISAV + isotype and ISAV + late α-PD-L1 (**Fig. 4A, Suppl. Datasheet**). For instance, early adjunct α-PD-L1 immunotherapy elicited significantly stronger dendritic-cell and M1-macrophage signaling than both ISAV + isotype and ISAV + late adjunct α-PD-L1, along with stronger activation of key cytokine pathways (e.g., IL-1, IL-2, IL-12) (**Fig. 4A-B, Fig. S6**). Likewise, early adjunct α-PD-L1 therapy significantly restored Th1 and Th17 signaling (**Fig. 4A-B, Fig. S6**). Importantly, our data revealed a strong (inverse) overlap between immune pathways affected by IPM-induced immune paralysis and those induced or restored by early adjunct checkpoint blockade (**Fig. 4A, Suppl. Datasheet**).

**Figure 4.**
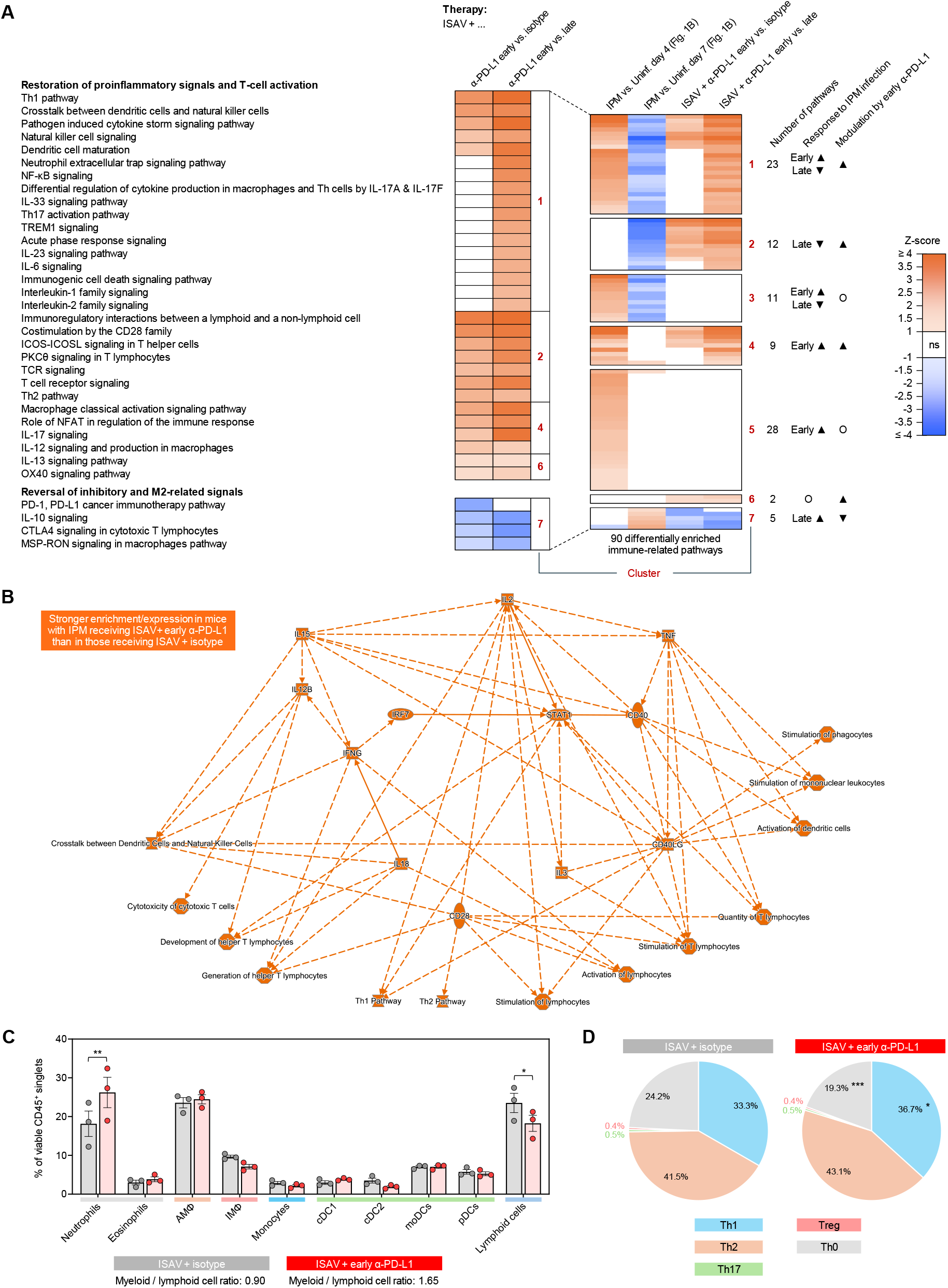
Early multimodal therapy with ISAV and adjunct α-PD-L1 attenuates exhaustion signals and enhances antifungal immune defense. (**A**) nCounter-based canonical pathway enrichment analyses performed on lung tissue homogenates of CPM-immunosuppressed mice with IPM on day 10 post-infection. The heatmap compares enrichment of selected pathways in mice that received ISAV + early adjunct α-PD-L1 versus ISAV + isotype (left column) and ISAV + late adjunct α-PD-L1 (right column). Orange and blue color indicate significantly stronger and weaker enrichment in the ISAV + early adjunct α-PD-L1 group compared to the reference groups, respectively. Pathways were selected from a total of 51 immune-related pathways with differential enrichment in response to early adjunct α-PD-L1 group (clusters 1, 2, 4, 6, and 7 in the insert). N = 3 mice per treatment arm. See **Suppl. Datasheet** for complete data. (**B**) Network of changes to the pulmonary immune transcriptome in mice with IPM that received ISAV + early adjunct α-PD-L1 immunotherapy versus ISAV + isotype, as predicted by Ingenuity Pathway Analysis (day 10 post-infection). N = 3 mice per treatment arm. (**C**) Flow cytometric characterization of leukocyte subsets in lung tissue homogenates of mice with IPM according to the treatment arm (day 10 post-infection). Individual datapoints, means, and standard errors of the mean are shown. Colored bars below the x-axis refer to the overarching cell populations shown in **Fig. S7**. N = 3 mice per treatment arm. (**D**) Flow cytometric analysis of CD4^+^ T-helper cell subpopulations in mice with IPM according to the treatment arm. Means from 4-6 mice per treatment arm are shown. (**C-D**) Unpaired two-sided t-test. * p<0.05, ** p<0.01, *** p<0.001. Abbreviations (not including gene symbols in panel **B**): α = anti, AMΦ = alveolar macrophages, CD = cluster of differentiation, cDC = conventional dendritic cells, CTLA-4 = cytotoxic T lymphocyte-associated protein 4, ICOS(L) = inducible T-cell co-stimulator (ligand), IL = interleukin, IMΦ = interstitial macrophages, IPM = invasive pulmonary mucormycosis, ISAV = isavuconazonium sulfate, moDCs = monocyte-derived dendritic cells, MSP-RON = macrophage stimulating protein & *recepteur d’origine nantais*, NFAT = nuclear factor of activated T cells, NF-κB = nuclear factor kappa B, ns = not significant, OX40 = costimulatory receptor OX40 (CD134), PD-1 = programmed cell death protein 1, pDCs = plasmacytoid dendritic cells, PD-L1 = programmed death ligand 1, PKC = protein kinase C, TCR = T-cell receptor, Th = T-helper (cell), Treg = regulatory T cells, TREM = triggering receptors expressed on myeloid cells.

To further dissect the impact of early adjunct α-PD-L1 immunotherapy versus ISAV alone (plus isotype) on the pulmonary immune environment and effector cell accumulation, we performed flow cytometry on lung tissue homogenates on day 10 post-infection. Consistent with our previous observation in an IPA model^17^, adjunct α-PD-L1 immunotherapy led to a significantly elevated myeloid-to-lymphoid cell ratio in lung tissue (1.65 vs. 0.90, p=0.037, **Fig. 4C, Fig. S7**). Specifically, mice that received early ISAV + adjunct α-PD-L1 showed increased pulmonary accumulation of neutrophils compared to the ISAV + isotype group (26.2% vs. 18.2% of viable leukocytes, p=0.003, **Fig. 4C**).

We further profiled B and T cells in lung tissue homogenates on day 10 post-infection to assess differences in adaptive immune cell phenotypes. Mice receiving early adjunct α-PD-L1 showed a modest shift from naïve to differentiated T cells compared to those receiving ISAV + isotype (p<0.001, **Fig. 4D**). While the Th2 phenotype accounted for the largest proportion of Th cells in either treatment arm, the ISAV + early adjunct α-PD-L1 group showed a slight increase in Th1 cells compared to mice receiving ISAV + isotype (36.7% vs. 33.3%, p=0.046, **Fig. 4D**). In contrast, B-cell phenotypes, including the proportions of activated B cells, were similar in both treatment arms (**Fig. S8**).

Altogether, these findings suggest that early adjunct anti-PD-L1 therapy largely alleviated IPM-induced exhaustion signals, led to increased accumulation of neutrophils in the lung, and broadly stimulated T-cell differentiation, activation, and signaling (**Fig. 5**). The significant inverse overlap between suppressive immune signals in untreated mice with IPM and surrogates of α-PD-L1-induced immune augmentation, combined with a lack of local or systemic immunotoxicity, suggests largely specific attenuation of infection-induced immune paralysis by early adjunct ICI therapy.

**Figure 5.**
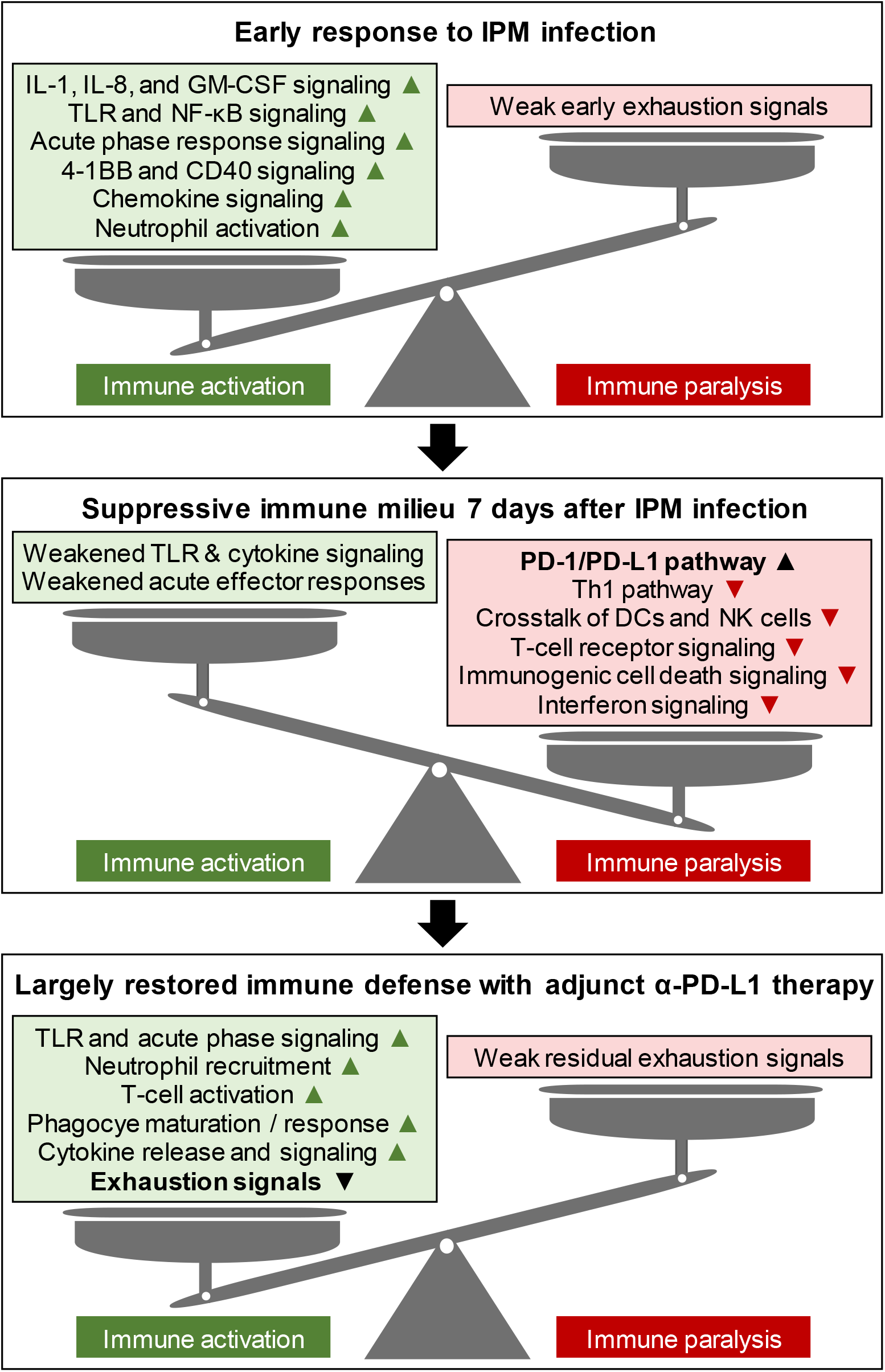
Schematic model summarizing IPM-induced immune paralysis signals and their alleviation by early adjunct α-PD-L1 immunotherapy. Abbreviations: 4-1BB = costimulatory receptor 4-1BB (CD137), α = anti, CD = cluster of differentiation, DCs = dendritic cells, IL = interleukin, IPM = invasive pulmonary mucormycosis, NF-κB = nuclear factor kappa B, NK cells = natural killer cells, PD-1 = programmed cell death protein 1, PD-L1 = programmed death ligand 1, Th = T-helper (cell), TLR = Toll-like receptor.

## Discussion

Several preclinical studies and case reports suggested ICI therapy as an effective and rapidly available (non-cellular) adjunct immunotherapy in hosts with IFDs^7^. To better understand the optimal “window of opportunity” for ICI therapy, we explored the early dynamics of immune paralysis in mice with IPM and other mold pneumonias. Despite the use of only neutrophil-targeted chemotherapy without the confounding effect of corticosteroids on immune exhaustion pathways^18,23^ and relatively low fungal inoculums, we found a rapid shift of the pulmonary immune environment from initial inflammation to severe immune paralysis within one week after infection.

Notably, our focus on pathway-level analyses allowed us to determine the “net state” of antifungal immunity in a complex and dynamically evolving immune environment with a plethora of feedback loops and intricate relationships^24^. For instance, transcription of the CTLA-4 gene itself was (already) suppressed by day 7 post-infection, whereas CTLA-4 and PD-1/PD-L1 checkpoint signaling was significantly induced on pathway level. This observation was consistent with both the overall tolerogenic immune state in our mouse model and a recent report of rapid induction of immune checkpoint signaling in mucormycosis patients^25^. Additionally, we found strongly diminished cytotoxic responses such as granzyme production and NK-cell signaling. While the interplay of Mucorales with cytotoxic T cells remains poorly understood, our data aligns with a prior report of suppressed NK-cell activity after co-culture with *Rhizopus* hyphae *in vitro*^26^.

The relative contributions of underlying host immune compromise, tissue damage and repair, host immune feedback loops, the infection microenvironment, and specific fungal virulence factors to immune paralysis remain largely unknown^7,25^. Although direct interaction of fungal ligands with checkpoint receptors have been described for pathogenic yeasts^27,28^, our observation of largely overlapping immune paralysis signatures in response to three evolutionarily and biologically vastly dissimilar molds^29^ points to a predominance of host-mediated drivers such as tissue inflammation. Whether known Mucorales virulence factors involved in early host interactions (e.g., the fungal spore coat protein CotH3^30^) and tissue damage (e.g., the mucoricin toxin^31^) contribute to immune paralysis requires further study.

Effective post-infection ICI therapy is thought to rely on a delicate balance between reversibility of immune exhaustion (e.g., residual self-renewal capacity of T cells) and antigen burden, implying that timing of immunotherapy may be critical^32^. In clinical mycology, immunomodulators are often used as a (late-stage) salvage intervention in patients with intractable infections. In contrast, preclinical studies of ICIs and other immunomodulators in fungal infection models have mostly focused on very early immunotherapy, often without concomitant antifungals^7,15,18,19^. This conceptual difference has likely been a major contributor to the current “bench-bedside disconnect” of antifungal immunotherapy^7^. Focusing on clinically viable timepoints, i.e., after infection-induced distress had manifested, we therefore compared the time-dependent benefits of ICIs given in the context of Mucorales-active antifungal therapy. Although not reaching significance due to the strong overall benefit of adjunct α-PD-L1 and limited power of non-parametric multi-group comparisons, we observed a trend toward stronger improvement in morbidity with early adjunct α-PD-L1 therapy. Furthermore, early adjunct α-PD-L1 therapy provided significantly stronger reinvigoration of antifungal immunity than sequential administration of ISAV and α-PD-L1. Reflecting the limited evidence for benefits of salvage immunotherapy in clinical reports^14,33^, this observation underscores the importance of studying early deployment of host-targeted therapy in high-risk patient populations with opportunistic mold infections.

Although the few published case reports of ICI therapy in clinical mycology had used nivolumab (α-PD-1)^11–14^, we opted for α-PD-L1 which yielded a stronger (mono-)therapeutic benefit and caused less IL-6-driven hyperinflammation than α-PD-1 in our previous study^18^. ICI dosing has been highly heterogenous in both clinical case reports and experimental mycology studies^7^. While dosing regimens are not immediately transferable between murine and human ICI agents, we and others previously reported improved tolerability and outcomes with sub-oncologic dosing of α-PD-(L)1 in fungal infection models^17,34^. Combining this approach with first-line antifungal therapy, we found relatively specific reversal of infection-induced immune paralysis signals without signs of immunotoxicity or systemic hyperinflammation. While these findings provide important safety signals to support clinical evaluation of ICIs as early antifungal immunotherapy, future studies should explore inoculum-immune exhaustion relationships and delineate the impact of ICI dosing on restoration of protective immune responses to maximize therapeutic benefits while avoiding immunotoxicity.

In addition to novel insights into pulmonary immune dynamics after IPM infection and α-PD-L1 therapy, our findings confirm several prior clinical and preclinical observations. For instance, our transcriptional data corroborated a strong relationship between impairment of mononuclear phagocyte activation/maturation and checkpoint induction, as suggested by clinical case reports^11,35^. Furthermore, our findings of predominantly Th2-polarized T-helper cells in the lung and robust IL-4 serum levels align with the strong Th2 activity observed in patients with hematologic malignancy and mucormycosis^36^. However, BALB/c mice are strongly Th2 biased^37^; therefore, these results would need to be confirmed in other genetic backgrounds. Additionally, our finding of a rather modest shift toward Th1 polarization in α-PD-L1-treated mice with IPM is consistent with prior reports of limited effects of ICI therapy on Th-cell phenotypes after invasive mold infections^7,17,18^. Therefore, systematic exploration of combination immunotherapy with ICIs and the Th1-polarizing cytokine IFN-γ, as described in some case reports^11,12,14,35^, may be warranted. Lastly, our data corroborated enhanced accumulation of neutrophils at the site of infection as a key component of ICI activity in boosting antifungal immunity^17^. Experimental data in a non-neutropenic candidiasis model showed increased neutrophil release from the bone marrow through PD-L1-dependent modulation of autocrine chemokine signals in precursor cells^38^. However, the detailed mechanisms of neutrophil recruitment in mold pneumonias affecting hosts prone to quantitative or functional neutropenia^2,4^ require further study.

This study has several limitations. Firstly, we used otherwise healthy mice with chemotherapy-induced neutropenia. Although common practice in experimental mycology, there is an evident need to confirm and compare ICI efficacy in models of other common mucormycosis manifestations (e.g., sino-orbital or disseminated disease) and additional host background such as uncontrolled diabetes, active hematologic malignancy, or trauma. Secondly, while our comparison of immune paralysis signatures in IPM, IPA, and IF focused on shared motifs, these analyses relied on a single fungal isolate per genus and did therefore not account for the significant inter-species and inter-isolate variation in immunopathology^39^. Thirdly, although we precluded a systemic cytokine storm and confirmed the absence of ICI-induced pneumonitis, we did not systematically screen other organs (e.g., the gastrointestinal tract) for signs of offsite immunotoxicities. Our study also did not account for long-term changes in the Mucorales-reactive and auto-reactive T-cell repertoire or other immunologic long-term effects after ICI therapy. Lastly, “alternative” checkpoint pathways that are druggable with investigational ICI agents (e.g., TIM-3, LAG-3, TIGIT)^40^ or other clinically important homeostatic pathways (e.g., angiogenesis) were insufficiently covered by our pathway-level analyses and would require detailed flow cytometric and tissue-level exploration in future studies.

In summary, we found rapidly emerging immune paralysis signals in mice with IPM and other mold pneumonias. IPM-associated immune paralysis was significantly attenuated upon early adjunct ICI therapy, supporting the concept of preemptive multimodal (antifungal and immunomodulatory) therapy in high-risk patients with mucormycosis. Given the paucity of immunologic data in human IPM patients, future analyses of well-annotated respiratory and blood-based patient specimens will be critical to confirm the immune dynamics observed in this study and finetune strategies for adjunct host-targeted therapies. Moreover, further improvement of early diagnosis^41^, identification of host biomarkers for early stratification of disease severity and immune status, novel first-in-class antifungals^42^, testing of other adjunct immunomodulators, and utilization of innovative trial designs will be pivotal to advance personalized management of IPM and other deadly mold pneumonias.

## Materials and Methods

### Animal studies

Eight-week-old female BALB/c mice were immunosuppressed with cyclophosphamide (Long Grove Pharmaceuticals; 250 mg/kg on day −4, 150 mg/kg on day −1, and 100 mg/kg twice weekly thereafter). On day 0, mice were infected intranasally with 50,000 spores of *Rhizopus arrhizus* (clinical isolate Ra-749), 50,000 conidia of *Aspergillus fumigatus* (reference strain Af-293), or 150,000 spores of *Fusarium solani* (clinical isolate Fs-001).

For therapeutic studies in the IPM model, treatment with ISAV was initiated on day 3 post-infection (64 mg/kg by oral gavage twice daily for 5 days), when the mice had developed significant infection-induced distress. Additionally, mice received 0.25 mg/kg α-PD-L1 (Leinco Technologies, #P371) or the corresponding IgG2b isotype antibody (Leinco Technologies, #I-1034) intraperitoneally either on days 3+5 (early) or days 6+8 (late).

Infection severity was scored using the modified murine sepsis score, as described previously^18,21^.

### Quantification of fungal burden

Total genomic DNA was isolated from murine lung tissue using the DNeasy Tissue Kit (Qiagen). Fungal burden was determined by quantitative polymerase chain reaction using primers and probes for the Mucorales 18S rRNA gene^43^ and was compared to a standard curve compiled using murine lung tissue specimens spiked with 10^2^-10^7^ *R. arrhizus* spores.

### Transcriptional analyses

Lungs were weighed and flash-frozen in 1 mL RNAprotect Tissue Reagent (Qiagen). For RNA isolation, lungs were thawed and homogenized in 1 mL RLT Buffer (Qiagen) using a Mini Bead-Beater (Biospec Products) and 3-mm glass beads (Sigma-Aldrich). Total RNA was isolated from 30 mg of homogenized tissue using the RNeasy Tissue Kit (Qiagen) and RNase-Free DNase Set (Qiagen) according to the manufacturer’s instructions. RNA concentration, purity, and integrity were analyzed using a NanoDrop OneC spectrophotometer (Thermo Fisher Scientific), Qubit 3.0 fluorometer (Thermo Fisher Scientific), and TapeStation 4200 (Agilent Technologies).

Expression of immune-related genes was assessed using the 785-plex nCounter Immune Exhaustion panel (NanoString Technologies) according to the manufacturer’s instructions. Data was analyzed using the nSolver Analysis Software (NanoString Technologies), with background thresholding to the median of negative controls and normalization to the panel’s 12 housekeeping genes (geometric mean). Genes with absolute inter-group fold changes >1.5 and p-values <0.05 were considered significantly differentially expressed.

Considering genes with fold expression levels of >|1.5|, pairwise mean inter-group expression ratios were imported into Ingenuity Pathway Analysis (Qiagen) for canonical pathway enrichment analysis^44^. Differences in canonical pathway enrichment were considered significant at an absolute z-score ≥1 and Benjamini-Hochberg adjusted p-value <0.05. The NanoString Cell Type Profiling Module was used to determine cell abundance scores based on known ratios of immune cell markers^45,46^.

### Flow cytometry

Lung tissue processing to obtain single-cell suspensions was performed according to published protocols^45^. For the myeloid and B-cell panels, samples were incubated for 30 min with fluorescent antibodies at a concentration of 1:250 and washed in PBS + 1% fetal bovine serum (FBS). For the T-cell panel, the Inside Stain Kit (Miltenyi Biotec) was used according to the manufacturer’s instructions. Antibody panels are summarized in **Supplementary Methods**. Prior to measurement, cells were resuspended in PBS + 1% FBS + 2% paraformaldehyde and passed through a cell strainer cap. Flow cytometry was performed on a Cytek Aurora Analyzer. Data was analyzed using FlowJo v10.8.1.

### Luminex assay

Serum specimens were cryopreserved following published protocols^17^. Cytokine and chemokine concentrations were determined using a 13-plex magnetic cytokine panel (LXSAMSM-13, R&D Systems) and Luminex 200 reader (Luminex Corporation) according to the manufacturers’ manuals.

### Statistical analyses

Significance testing was performed using the Mann-Whitney U test or unpaired t-test for two-group comparisons and one-way analysis of variance with Tukey’s post-test or Kruskal-Wallis test with Dunn’s post-test for multi-group comparisons, as appropriate. Significance tests are specified in the figure legends. R v4.3.1, Excel 365 (Microsoft Corporation), and Prism v10 (GraphPad Software) were used for data compilation, analysis, and visualization.

## Supporting information

Suppl. Datasheet

## Footnote Page

### Regulatory approval

This study involving murine infection models was reviewed and approved by the University of Texas MD Anderson Cancer Center Institutional Animal Care & Use Committee (protocol 00001734-RN02).

### Author contributions

SW and DPK designed the study and acquired funding. SW, JPG, YW, NDA, UB, and LM performed experiments. SW and JPG analyzed and visualized data. REL provided critical feedback and contributed to funding acquisition. SW, SEE, and DPK supervised the study. SW and DPK wrote the manuscript. JPG, YW, LM, REL, and SEE edited the manuscript. All authors approved the final version of the manuscript.

## Acknowledgements

This study was supported by an investigator sponsored study grant from Astellas Pharma Global Development (to DPK and SW). The sponsor had no influence on the research design, data analysis, and decision to publish the findings of this study. This study was further supported by an intramural Internal Medicine Research Award (to SW), the Cyrus Scholar Award (to SW), and the Robert C. Hickey Chair Endowment (to DPK). MD Anderson’s Advanced Technology Genomics Core was supported in part by intramural funding and Cancer Center Support Grant P30 CA016672. The authors thank Yun Zhu and Denaha J. Doss for their excellent support with nCounter analyses.

## Prior presentations

Preliminary results of this study have been presented at ESCMID Global 2024 (Barcelona, Spain) and IDWeek 2024 (Los Angeles, CA, United States).

