## Supplement for "Early adjunct anti-PD-L1 immunotherapy improves outcomes and restores infection-induced immune paralysis in mice with invasive pulmonary mucormycosis"

### Supplementary Materials

Supplementary Methods

Suppl. Table S1

Suppl. Figures S1-S8

**Supplementary Methods**

Histopathology

Representative mouse lung tissue sections were fixed and stained with hematoxylin & eosin following published protocols[S1]. Briefly, mouse lungs were fixed intratracheally and incubated for 24 hours with 10% formalin. Thereafter, lungs were transferred to 70% ethanol and embedded in paraffin. Formalin-fixed paraffin-embedded tissue blocks were cut into 10 µm sections, mounted, deparaffinized with xylene, washed with ethanol, and rehydrated. After hematoxylin & eosin staining, slides were imaged using a BZ-X810 microscope (Keyence) at 20-fold magnification.

Antibodies used for flow cytometry studies

| Myeloid panel | | |
|---|---|---|
| Materials / Resources | Reference / Identifier | Source |
| CD45 (30-F11) – RedFluor 710 | Cat # 80-0451-U100 | Tonbo Biosciences |
| CD11b (M1/70) – PerCP-Cy5.5 | Cat # 45-0112-82 | Invitrogen |
| CD11c (N418) – Brilliant Violet 737 | Cat # 569236 | BD Horizon |
| MHC II (M5/114.15.2) – PE-Cy 7 | Cat # 107630 | BioLegend |
| Ly-6 C (HK1.4) – Brilliant Violet 650 | Cat # 128049 | BioLegend |
| Ly-6 G (1A8) – PE | Cat # 127608 | BioLegend |
| CD64 (X54-5/7.1) – Brilliant Violet 711 | Cat # 139311 | BioLegend |
| Siglec-F (E50-2440) – PE-CF594 | Cat # 562757 | BD Biosciences |
| F4/80 Antigen (BM8) – FITC | Cat # 11-4801-82 | Invitrogen |
| CD103 (2E7) – Brilliant Violet 785 | Cat # 121439 | BioLegend |
| CCR2(SA203G11) – APC | Cat # 150628 | BioLegend |
| Ghost Dye UV 450 | Cat # 13-0868-T100 | Tonbo Biosciences |

| T-cell panel | | |
| --- | --- | --- |
| Materials / Resources | Reference / Identifier | Source |
| CD45 (30-F11) – RedFluor 710 | Cat # 80-0451-U100 | Tonbo Biosciences |
| CD3e (145-2C11) – PerCP-Cy 5.5 | Cat # 65-0031-U100 | Tonbo Biosciences |
| CD4 (RM4-5) – PE-Cy 7 | Cat # 60-0042-U100 | Tonbo Biosciences |
| CD8a (53.6.7) – APC-Cy 7 | Cat # 25-0081-U100 | Tonbo Biosciences |
| FOXP3 (MF-14) – Alexa Fluor 647 [#] | Cat # 126408 | BioLegend |
| Gata-3 (TWAJ) – Alexa Fluor 488 [#] | Cat # 53-9966-42 | Invitrogen |
| T-bet (4B10) – Brilliant Violet 785 [#] | Cat # 644835 | BioLegend |
| ROR γ T (AFKJS-9) – PE [#] | Cat # 12-6988-80 | Invitrogen |
| PD-1 (29F.1A12) – Brilliant Violet 421 | Cat # 135221 | BioLegend |
| Ghost Dye UV 450 | Cat # 13-0868-T100 | Tonbo Biosciences |

[#] Intracellular marker

| B-cell panel | | |
| --- | --- | --- |
| Materials / Resources | Reference / Identifier | Source |
| IgA (C10-1) – Brilliant Violet 395 | Cat # 743299 | BD Biosciences |
| CD27 (LG.3A10) – Brilliant Violet 496 | Cat # 741094 | BD Biosciences |
| CD43 (S7) – Brilliant Violet 563 | Cat # 741238 | BD Biosciences |
| CD3 (17A2) – Brilliant Violet 661 | Cat # 741562 | BD Biosciences |
| CD1d (1B1) – Brilliant Violet 737 | Cat # 741758 | BD Biosciences |
| CD5 (53-7.3) – Brilliant Violet 421 | Cat # 100618 | BioLegend |
| CD21 (7E9) – Brilliant Violet 510 | Cat # 123437 | BioLegend |
| CD138 (281-2) – Brilliant Violet 605 | Cat # 142516 | BioLegend |
| CD19 (6D5) – Brilliant Violet 711 | Cat # 115555 | BioLegend |
| CXCR5 (L138D7) – Brilliant Violet 785 | Cat # 145523 | BioLegend |
| IgM (RMM-1) – PE-Cy5 | Cat # 406544 | BioLegend |
| CD23 (B3B4) – PE-Cy7 | Cat # 101614 | BioLegend |
| B220 (RA3-6B2) – Alexa Fluor 700 | Cat # 103232 | BioLegend |
| IgD (11-26c.2a) – APC-Cy7 | Cat # 405716 | BioLegend |
| Ghost Dye UV 450 | Cat # 13-0868-T100 | Tonbo Biosciences |

### Supplementary Table

**Table S1.** Mean serum cytokine and chemokine levels (pg/mL) in mice with invasive pulmonary mucormycosis according to the treatment arm. Unpaired two-sided t-test. ND denotes non-detectable levels (below the lower limit of detection) in at least half of the mice.

| Analyte | Limit of detection | Day 7 post-infection | | | Day 10 post-infection | | |
|---|---|---|---|---|---|---|---|
| | | N = 3 | N = 3 | P-value | N = 4 | N = 4 | P-value |
| CCL2 | 38.9 | ND | ND | --- | ND | ND | --- |
| CCL3 | 0.1 | 0.5 | 0.4 | 0.557 | 0.6 | 0.6 | 0.547 |
| CCL4 | 18.2 | ND | ND | --- | ND | ND | --- |
| CXCL2 | 0.2 | 1.2 | 0.6 | 0.493 | 4.5 | 2.9 | 0.091 |
| GM-CSF | 1.0 | ND | ND | --- | ND | ND | --- |
| IFN-γ | 1.7 | ND | ND | --- | ND | ND | --- |
| IL-2 | 1.0 | 1.4 | ND | --- | 2.8 | 1.4 | 0.198 |
| IL-4 | 6.0 | 19.0 | 19.4 | 0.803 | 19.0 | 22.2 | 0.565 |
| IL-6 | 2.9 | ND | ND | --- | 5.1 | ND | --- |
| IL-12 p70 | 4.7 | ND | ND | --- | ND | ND | --- |
| IL-17A | 4.5 | ND | ND | --- | ND | ND | --- |
| IL-33 | 10.4 | ND | ND | --- | ND | ND | --- |
| TNF-α | 0.2 | ND | ND | --- | ND | ND | --- |

ISAV + isotype    ISAV + α-PD-L1

### Supplementary Figures

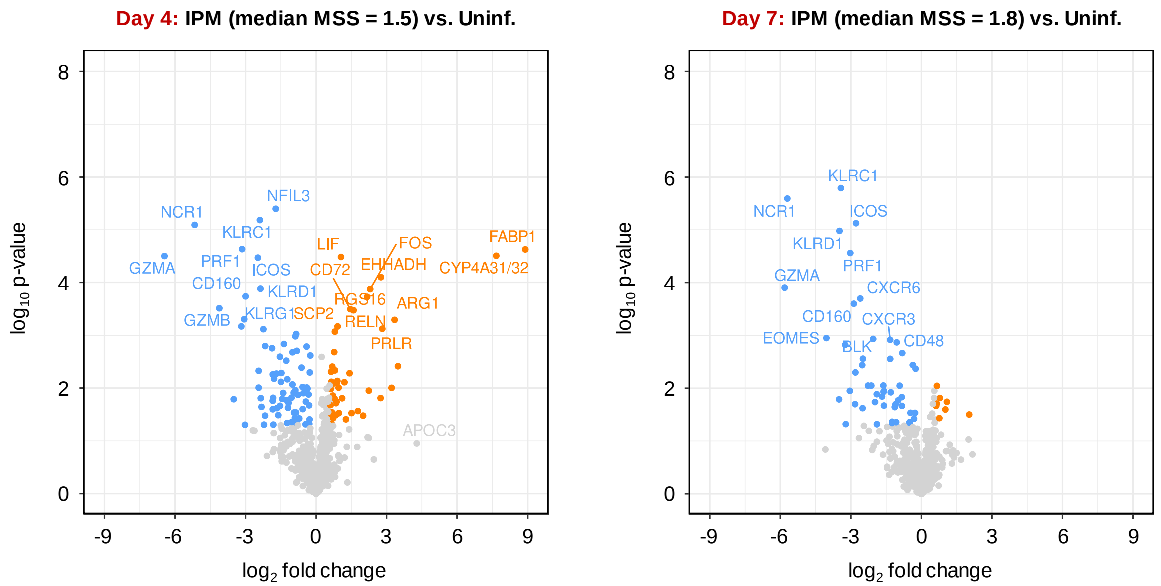


**Figure S1.** Volcano plots comparing gene expression levels in lung tissue of mice with IPM versus uninfected mice on days 4 and 7 post-infection using the 785-gene nCounter Immune Exhaustion panel. Significantly differentially expressed genes (absolute fold change >1.5, $p<0.05$) are color-coded, with orange and blue color indicating upregulated and downregulated expression in mice with IPM, respectively. The right panel is identical to the IPM panel in **Fig. 2A**. N = 3 mice per group.

<u>Abbreviations (not including individual gene symbols in the plots):</u> IPM = invasive pulmonary mucormycosis, MSS = (modified) murine sepsis score, Uninf. = uninfected.

**A**

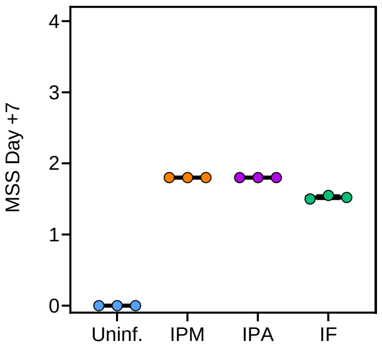


**B** **Top 10 downregulated immune-related canonical pathways by day 7 post-infection**

| **Invasive pulmonary mucormycosis** | Z-score |
|---|---|
| Th1 pathway | -4.24 |
| Immunoregulatory interactions between a lymphoid and a non-lymphoid cell | -4.03 |
| Costimulation by the CD28 family | -3.32 |
| TCR signaling | -3.16 |
| ICOS-ICOSL signaling in T helper cells | -3.16 |
| PKCθ signaling in T lymphocytes | -3.16 |
| Communication between innate and adaptive immune cells | -3.00 |
| T cell receptor signaling | -2.68 |
| Differential regulation of cytokine production in macrophages and T-helper cells by IL-17A and IL-17F | -2.65 |
| TREM1 signaling | -2.45 |

| **Invasive pulmonary aspergillosis** | Z-score |
|---|---|
| Th1 pathway | -5.20 |
| Immunoregulatory interactions between a lymphoid and a non-lymphoid cell | -5.20 |
| Costimulation by the CD28 family | -3.87 |
| TCR signaling | -3.64 |
| Role of NFAT in regulation of the immune response | -3.41 |
| Interferon gamma signaling | -3.36 |
| Immunogenic cell death signaling pathway | -3.32 |
| ICOS-ICOSL signaling in T helper cells | -3.15 |
| Pathogen induced cytokine storm signaling pathway | -3.12 |
| Neutrophil degranulation | -3.05 |

| **Invasive fusariosis** | Z-score |
|---|---|
| TCR signaling | -4.00 |
| Immunoregulatory interactions between a lymphoid and a non-lymphoid cell | -3.96 |
| Costimulation by the CD28 family | -3.36 |
| Neutrophil degranulation | -3.05 |
| Pathogen induced cytokine storm signaling pathway | -3.02 |
| Th1 pathway | -2.75 |
| Neutrophil extracellular trap signaling pathway | -2.67 |
| Crosstalk between dendritic cells and natural killer cells | -2.67 |
| Interleukin-1 family signaling | -2.50 |
| Communication between innate and adaptive immune cells | -2.32 |

**Figure S2.** (**A**) MSS scores on day 7 post-infection according to the pathogen challenge received. (**B**) Top 10 downregulated canonical pathways in mice with mold pneumonias (day 7 post-infection) compared to uninfected cyclophosphamide-immunosuppressed mice. N = 3 mice per group.

Abbreviations: CD = cluster of differentiation, ICOS(L) = inducible T-cell co-stimulator (ligand), IL = interleukin, IF = invasive fusariosis, IPA = invasive pulmonary aspergillosis, IPM = invasive pulmonary mucormycosis, MSS = (modified) murine sepsis score, NFAT = nuclear factor of activated T cells, PKC = protein kinase C, TCR = T-cell receptor, Th = T-helper (cell), TREM = triggering receptors expressed on myeloid cells, Uninf. = uninfected.

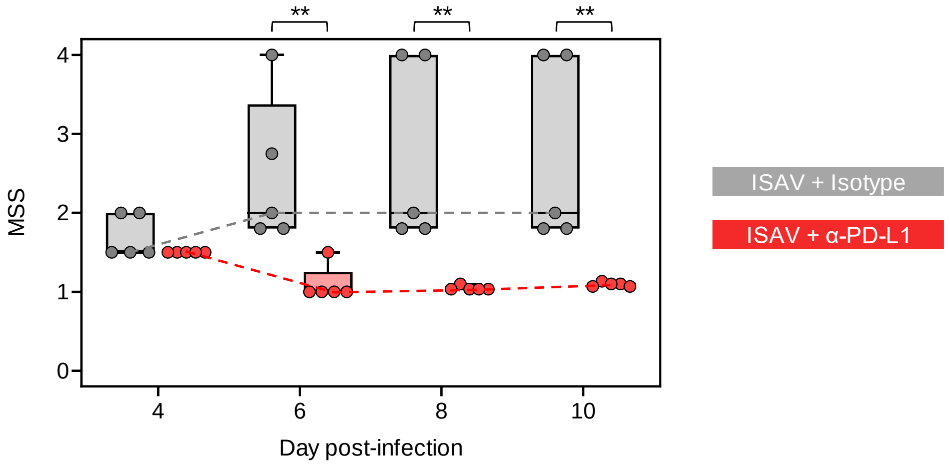


**Figure S3.** Evolution of MSS scores in mice with IPM according to the treatment arm. N = 5 mice per group from one representative experiment. Individual data points, quartiles (boxes), and ranges (error bars) are shown. Significance testing for each timepoint was performed using the Mann-Whitney U test. ** $p<0.01$. Of note, RNA sampling for analyses presented in **Fig. 4A-B** was performed on lung tissue from non-moribund survivors of this experiment (n = 3 per group) after MSS scoring on day 10.

Abbreviations: α = anti, IPM = invasive pulmonary mucormycosis, ISAV = isavuconazonium sulfate, MSS = (modified) murine sepsis score, PD-L1 = programmed death ligand 1.

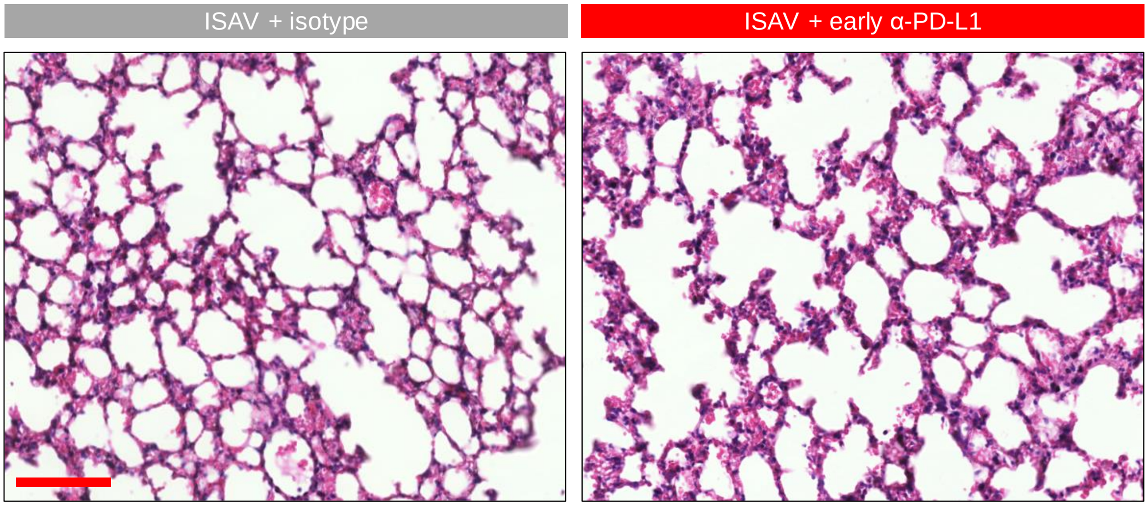


**Figure S4.** Hematoxylin & eosin-stained lung tissue sections according to the treatment arm, showing an absence of lymphocytic infiltrates, even in mice that received (early) adjunct α-PD-L1. Scale: 100 μm.

Abbreviations: α = anti, ISAV = isavuconazonium sulfate, MSS = (modified) murine sepsis score, PD-L1 = programmed death ligand 1.

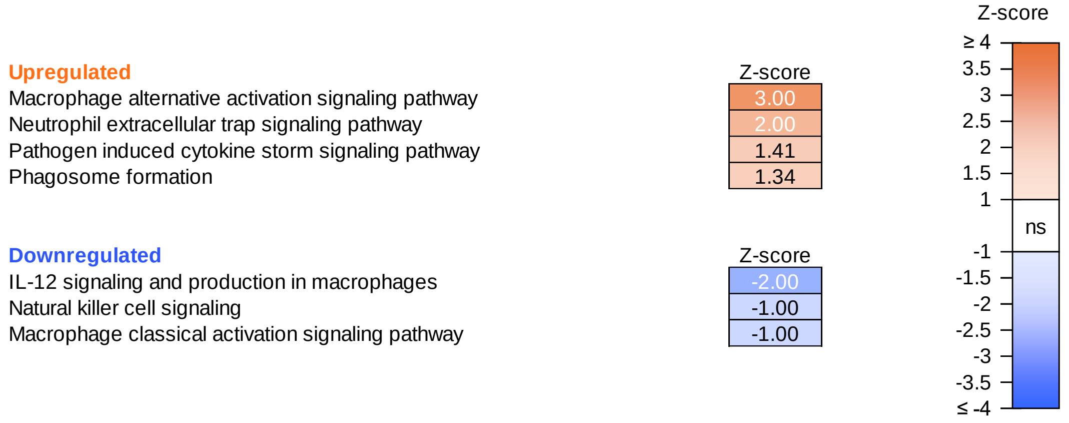


**Figure S5.** Differentially enriched canonical pathways in mice with invasive pulmonary mucormycosis that received antifungal therapy with isavuconazonium sulfate (+ isotype control) compared to uninfected immunosuppressed mice. Day 10 post-infection. N = 3 mice per group.

Abbreviations: IL = interleukin, ns = not significant.

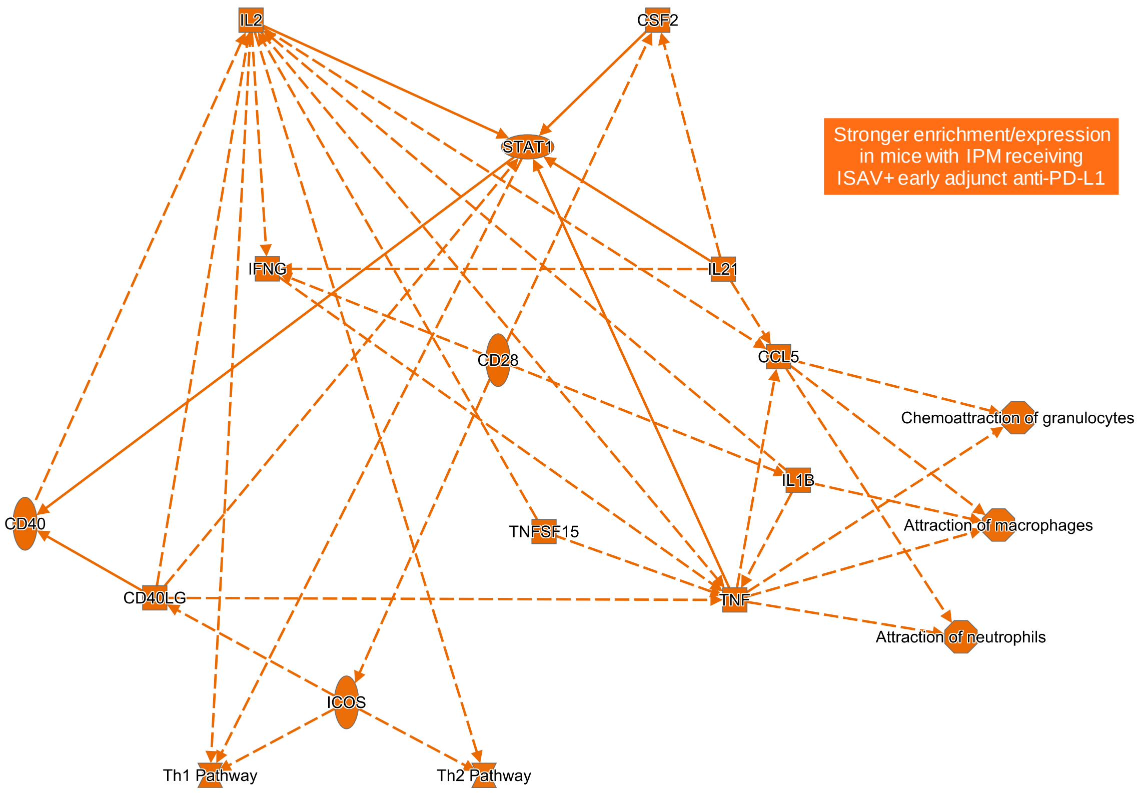


**Figure S6.** Network of differences in the pulmonary immune environment in mice with IPM that received ISAV plus early versus those receiving late adjunct α-PD-L1, as predicted by Ingenuity Pathway Analysis. N = 3 mice per group. Solid lines = direct interaction, dashed lines = indirect interaction.

Abbreviations (not including individual gene symbols in the network): α = anti, IPM = invasive pulmonary mucormycosis, ISAV = isavuconazonium sulfate, PD-L1 = programmed death ligand 1.

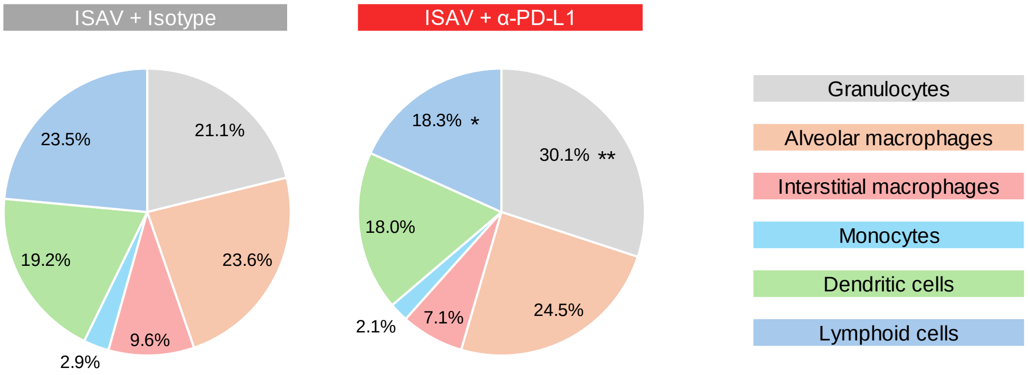


**Figure S7.** Flow cytometric analysis of leukocyte subsets (normalized to $CD45^+$ viable singlets) in mice with invasive pulmonary mucormycosis according to the treatment arm. Means from 3 mice per group are shown. Unpaired two-sided t-test. * $p<0.05$, ** $p<0.01$.

<u>Abbreviations:</u> α = anti, ISAV = isavuconazonium sulfate, PD-L1 = programmed death ligand 1.

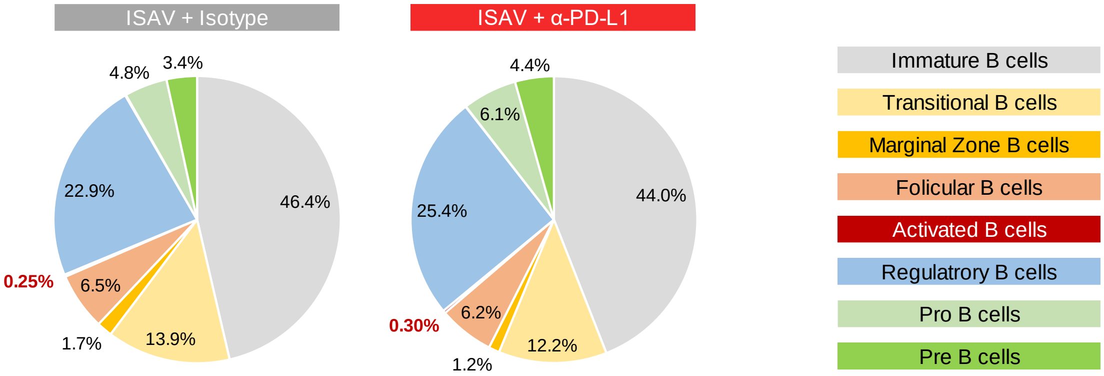


**Figure S8.** Flow cytometric analysis of B-cell phenotypes (normalized to $CD19^+$ viable singlets) in mice with invasive pulmonary mucormycosis according to the treatment arm. N = 3 mice per group.

<u>Abbreviations:</u> α = anti, ISAV = isavuconazonium sulfate, PD-L1 = programmed death ligand 1.
